# Predicting X-ray Diffuse Scattering from Translation Libration Screw Structural Ensembles

**DOI:** 10.1101/012955

**Authors:** Andrew H. Van Benschoten, Pavel V. Afonine, Thomas C. Terwilliger, Michael E. Wall, Colin J. Jackson, Nicholas K. Sauter, Paul D. Adams, Alexandre Urzhumtsev, James S. Fraser

## Abstract

Identifying the intramolecular motions of proteins and nucleic acids is a major challenge in macromolecular X-ray crystallography. While Bragg diffraction describes the average positional distribution of crystalline atoms, many different models can fit this distribution equally well. Diffuse X-ray scattering can reduce this degeneracy by directly reporting on correlated atomic displacements. Although recent technological advances are increasing the potential to accurately measure diffuse scattering, computational modeling and validation tools are still needed to quantify the agreement between experimental data and different parameterizations of crystalline disorder. A new tool, *phenix.diffuse*, addresses this need by employing Guinier’s equation to calculate diffuse scattering from Protein Data Bank (PDB)-formatted structural ensembles. As an example case, *phenix.diffuse* is applied to Translation-Libration-Screw (TLS) refinement, which models rigid body displacement for segments of the macromolecule. To enable calculation of diffuse scattering from TLS refined structures, *phenix.tls_models* builds multi-model PDB files that sample the underlying T, L and S tensors. In the glycerophosphodiesterase GpdQ, alternative TLS group partitioning and different motional correlations between groups yield markedly dissimilar diffuse scattering maps with distinct implications for molecular mechanism and allostery. These methods demonstrate how X-ray diffuse scattering can extend macromolecular structural refinement, validation, and analysis.

**Synopsis:** A method of simulating X-ray diffuse scattering from multi-model PDB files is presented. Despite similar agreement with Bragg data, different Translation-Libration-Screw refinement strategies produce unique diffuse intensity patterns.

## 1. Introduction

Protein flexibility is essential for enzymatic turnover, signaling regulation and protein-protein interactions (Fraser & Jackson, 2011). The motions enabling these functions span length-scales from a few angstroms to many nanometers and include transitions between side chain rotamers (Fraser *et al.*, 2009), loop openings and closings (Qin *et al.*, 1998; Williams *et al.*, 2014) and rigid-body subunit rotations (Korostelev & Noller, 2007). Multiple crystal structures are routinely compared to identify these motions and to derive hypotheses about the role of correlated motions in executing protein function. However, if only a single crystal form is available, evidence of concerted motion must be extracted from the spread in the electron density.

Extracting this information is possible because protein conformational heterogeneity across unit cells in space and within unit cells during the X-ray exposure time leads to an ensemble-averaged electron density map. Small atomic vibrations are commonly fit with individual B-factors, which describe the electron density distribution as a continuous isotropic Gaussian envelope around a central location and predominantly encompass disorder from thermal motion. Discrete conformational heterogeneity and crystal packing defects can be described as ensembles of structural models with partial occupancy (Burnley *et al.*, 2013; Rader & Agard, 1997; Gros *et al.*, 1990; van den Bedem *et al.*, 2009; Levin *et al.*, 2007; Wall *et al.*, 1997). If high-resolution diffraction data are available, anisotropic directionality can be added to B-factors by modeling a Gaussian distribution along each real-space axis, yielding an ellipsoid that shows the predominant direction of the electron density.

However, the large number of parameters required for anisotropic B-factor refinement renders it inaccessible for most macromolecular diffraction experiments. Translation-Libration-Screw (TLS) modeling, introduced by Schomaker and Trueblood (1968), can describe concerted, rigid-body displacement for groups of atoms (for comprehensive review see Urzhumtsev *et al.*, 2013). In TLS refinement, the target protein is segmented into independent rigid bodies that undergo small translations (“vibrations”) and rotations (“librations”). The anisotropic displacement of TLS refinement can be fully described with twenty parameters per rigid body, which can potentially contain many atoms. This small number of parameters compares favorably to the six parameters per atom demanded by individual anisotropic B-factor refinement and allows grouped anisotropic B-factors to be modeled at mid to low-resolution ranges. TLS refinement often leads to better agreement between observed and calculated structure factors, as measured by decreasing R_free_ values. The potential for improved statistics when relatively few observations are available has positioned TLS as a general refinement technique: 22% of the structures deposited in the Protein Data Bank (PDB; Bernstein *et al.*, 1977; Berman et al, 2000) employ TLS refinement in some form. TLS refinement is a component of many major structural refinement programs such as *Refmac* (Murshudov et al, 1997; Winn *et al.*, 2001), BUSTER-TNT (Bricogne et al, 1993, 2011) and *phenix.refine* (Afonine et al, 2012). These programs can select TLS groups automatically, based on biochemical intuition, or with the assistance of external web servers (Painter & Merritt, 2006a; Painter & Merritt, 2006b).

TLS refinement naturally suggests concerted structural motions, which can be assigned biological significance and subsequently tested with additional experiments. Visualization programs such as TLSViewer (Painter & Merritt, 2005) can convert the T, L and S tensors into a description of domain-scale mechanical motions and molecular graphics programs, such as Chimera (Pettersen et.al, 2004) or PyMol (DeLano, 2002), can be used to visualize the resulting anisotropic ellipsoids. For example, TLS refinement of the large multi-protein complex GroEL revealed subunit tilting that may play a role in transmitting conformational changes upon GroES or nucleotide binding (Chaudhry *et al.*, 2004) **(Figure 1a-b)**. Similarly, TLS modeling of the ribosome structure implied a “ratcheting” rotation of the 50S and 30S subunits around the peptidyl transferase center during tRNA translocation (Korostelev & Noller, 2007).

**Figure 1).**
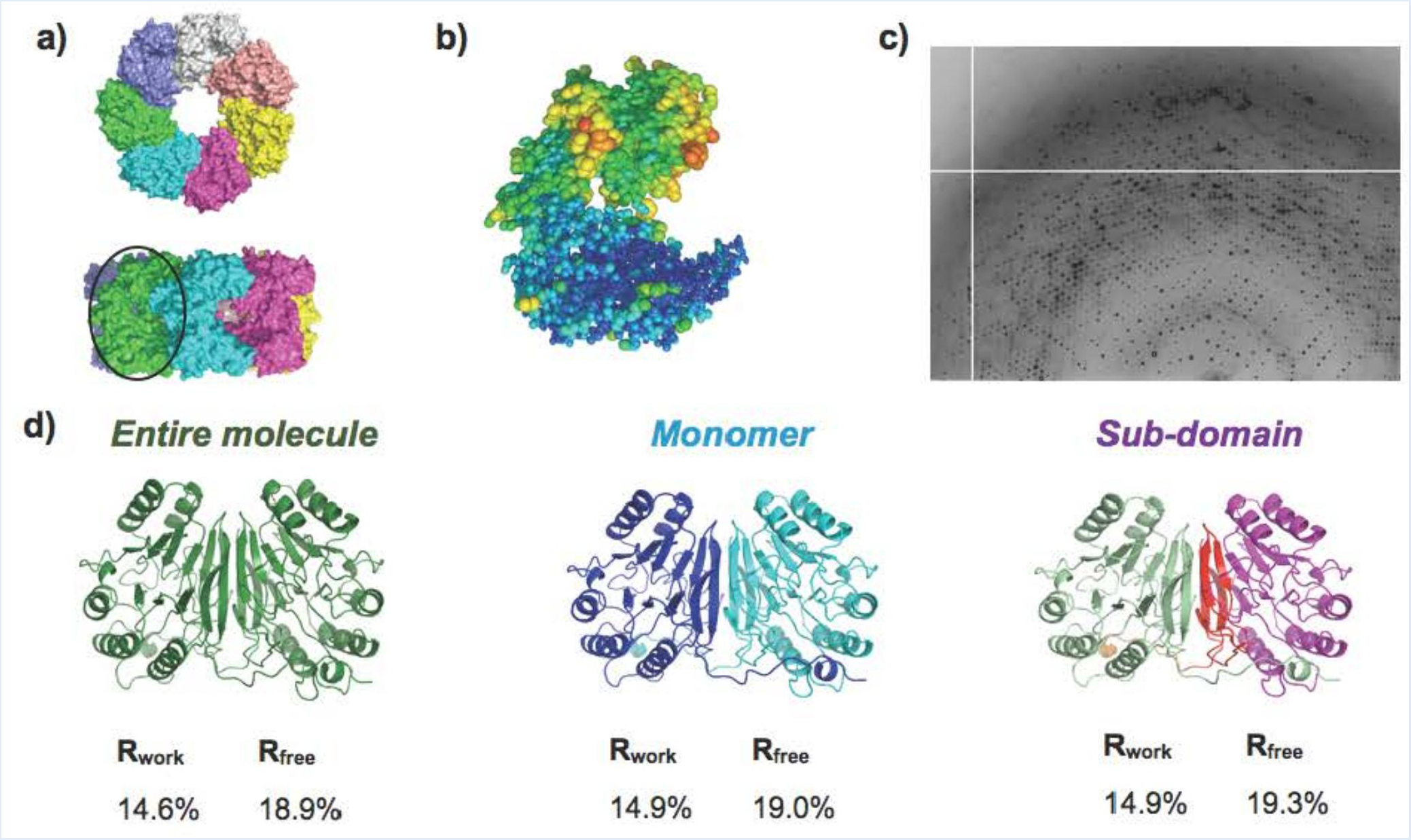
TLS refinement suggests macromolecular motions linked to function. **a)** Top and side view of GroEL. Each color denotes a unique chain. **b)** TLS refinement of GroEL subunits reveals a “tilting” motion around the center of the subunit. **c)** GpdQ diffraction image showing significant diffuse scattering features. **d)** Refinement of GpdQ fails to produce substantial changes in R_work_ and R_free_ values between alternate TLS groups. TLS refinement significantly improves the overall R_free_ (23.1% pre-TLS).

A potential complication of TLS refinement is that many different rigid body groupings can result in equivalent improvement of refinement statistics (Moore, 2009; Tickle & Moss, 1999). The inability to discriminate among alternate TLS models stems from the exclusive usage of Bragg diffraction data in model refinement. Because Bragg data reports on electron density averaged across all unit cells, there may be several models of structural displacement that fit the density equally well. Thus, TLS refinement might improve the modeled electron density but incorrectly describe the correlated motion that occurs in the crystal during the diffraction experiment. Drawing on additional sources of information such as patterns of steric clashes (van den Bedem *et al.*, 2013), NMR spectroscopy (Ruschak & Kay, 2012), or mutational analysis (Fraser *et al.*, 2009) can be used to distinguish competing models of correlated motion between non-bonded atoms.

An additional, yet rarely used, data source that can discriminate between these models is X-ray diffuse scattering from protein crystals, which results from correlated variation in the electron density distributions (Phillips *et al.*, 1980; Chacko & Phillips, 1992; Faure *et al.*, 1994; Clarage & Phillips, 1997; Mizuguchi *et al.*, 1994). Any deviation from a perfect crystal, such as thermal motion, crystal defects or static disorder breaks Bragg’s Law by violating the repetitive structure of the lattice, leading to diffraction in areas of reciprocal space outside of the fringe function. The theoretical relationship between conformational heterogeneity within unit cells and diffuse scattering has been available for decades (Guinier, 1963; Amorós & Amorós, 1968) and small-molecule crystallographers have used diffuse scattering data in refinement and model validation (Estermann & Steurer, 1998; Michels-Clark *et al.*, 2013, Welberry & Butler, 1994).

The potential of macromolecular diffuse scattering to break the degeneracy within refinement methods such as TLS, including information about the location and length scale of macromolecular disorder, has long been recognized (Thune & Badger, 1995; Perez *et al.*, 1996; Hery *et al.*, 1998; Tickle & Moss, 1999). Diffuse scattering maps predicted by models of motion can be calculated using either an all-atom covariance matrix or Guinier’s equation (Micu & Smith, 1994; Lindner & Smith, 2012). The covariance matrix describes correlated displacements between every pair of atoms, whereas Guinier’s equation models diffuse scattering from an ensemble of structure factors. Calculation of the covariance matrix has been used to model crystalline normal modes and TLS parameterization (Riccardi *et al.*, 2010). It is also possible to explicitly estimate each matrix element from molecular dynamics trajectories (Meinhold & Smith, 2007). The size of the covariance matrix scales as the square of the number of atoms, making full matrix calculations expensive to compute for large systems. This poses a significant challenge to quantitative diffuse scattering analysis. For these reasons, a straightforward method that calculates diffuse scatter from discrete multi-model PDB files may be preferable.

To meet this need, we developed *phenix.diffuse*, a new tool within the *Phenix* software suite (Adams *et al.*, 2010), which uses Guinier’s equation to calculate diffuse scattering from multi-model (ensemble) PDB files. Thus, *phenix.diffuse* can be applied to any motional model represented as an explicit ensemble of related structures. As a first application we have simulated the diffuse scattering produced by alternative TLS refinements of the glycerophosphodiesterase GpdQ (Jackson *et al.*, 2007). GpdQ is found in *Enterobacter aerogenes* and contributes to the homeostasis of the cell membrane by hydrolyzing the 3’-5’ phosphodiester bond in glycerophosphodiesters. Each chain of the dimeric enzyme contains three distinct structural elements: an α/β sandwich fold containing the active site, a domain-swapped active site cap and a novel dimerization domain comprised of dual-stranded antiparallel β-sheets connected by a small β-sheet. Though the catalytic mechanism of GpdQ is similar to other metallo-phosphoesterases, some substrates are too large to pass through the active site entrance as it is modeled in the crystal structure. Protein dynamics must therefore play a role in substrate entry and product release. Normal mode analysis of the GpdQ hexamer suggested high mobility in the cap domain and a breathing motion centered on the catalytic and dimerization domains (Jackson *et al.*, 2007). Due to the high global B-factors and presence of diffuse signal in the diffraction images **(Figure 1c)**, Jackson and colleagues performed three separate TLS refinements to model the crystalline disorder. All three TLS refinements improved the R_free_ values when compared to the standard isotropic B-factor refinement; however, there was no significant difference among the final R_free_ values from the refinements initiated with distinct TLS groupings. In contrast, our results reveal significant differences between the diffuse signals produced by each TLS refinement, highlighting the potential usefulness of diffuse scattering in optimizing structure refinement.

## 2. Methods

### 2.1. GpdQ refinement.

Based on the original refinement strategy of Jackson *et al.* (2007), we performed three different TLS refinements on the zinc-bound structure of GpdQ (PDB ID: 2DXN): *“Entire molecule”,* one TLS group for all residues; *“Monomer”*, one TLS group for each of the two individual chains; and *“Sub-domain”*, one TLS group for each of the α/β sandwich domain (residues 1-196), the “dimerization” domain (residues 197-255) and the “cap” domain (residues 257-271) of each chain. The pre-TLS refinement R_work_ and R_free_ were 19.1% and 23.1%, respectively. After defining the TLS groups, each structure was re-refined for 5 macrocycles in *Phenix.refine*. The strategy included refinement of the individual coordinates and isotropic B-factors, water picking and refinement of TLS parameters for defined TLS groups. Both the X-ray/atomic displacement parameters and X-ray/stereochemistry weights were optimized (Afonine *et al*, 2011). The final R_work_, R_free_ values for each refinement were as follows: *“Entire molecule”* (14.6%, 18.9%), *“Monomer”* (14.9%, 19.0%), *“Sub-domain”* (14.9%, 19.3%), suggesting approximately equal agreement with the Bragg data **(Figure 1d)**.

In TLS refinement, the eigenvalues of the T and L matrices describe the variance of the motional displacement along each orthogonal real-space axis. To avoid an unphysical description of TLS motion *(Urzhumtsev *et al.* accompanying manuscript)*, we inspected the eigenvalues of each TLS refinement to ensure non-negative eigenvalues for the T and L matrices **(Supplemental Table#1)**. Although solvent is expected to contribute significantly to experimental diffuse scattering, we removed water molecules after refinement. This step ensures that all subsequent diffuse scattering simulations only reflect correlated motions implicit in the TLS refinement.

### 2.2 phenix.tls_models and TLS ensemble generation.

We used *phenix.tls_models* (*Urzhumtsevet al., accompanying manuscript*) to convert the TLS matrices to a structural ensemble. *phenix.tls_models* receives as input a structure with TLS header information, separates the molecule into individual TLS groups and randomly samples the real-space distribution for each group based on mathematical decomposition of the T, L and S matrices. The sampled PDB files are then either re-assembled into a multi-model PDB ensemble or output with no further changes **(Figure 2)**. To ensure adequate sampling of the underlying Gaussian distributions, we generated ensembles of different sizes and monitored the convergence of the global correlation coefficient between diffuse maps in which spherically-symmetric sources of diffuse scattering have been removed (“anisotropic maps”: **Supplemental Table #2**). These maps offer an improved comparison relative to the raw diffuse signal because they correct for the resolution dependency of diffuse scattering, which would otherwise lead to overestimation of inter-map correlation. We determined that an ensemble size of 1000 models was sufficient for effective sampling of each TLS refinement. The extent of the motions predicted by the “sub-domain” refinement **(Supplemental Figure #1)** is quite surprising and likely results from a lack chemical restraints within the TLS refinement implementation in Phenix.

**Figure 2).**
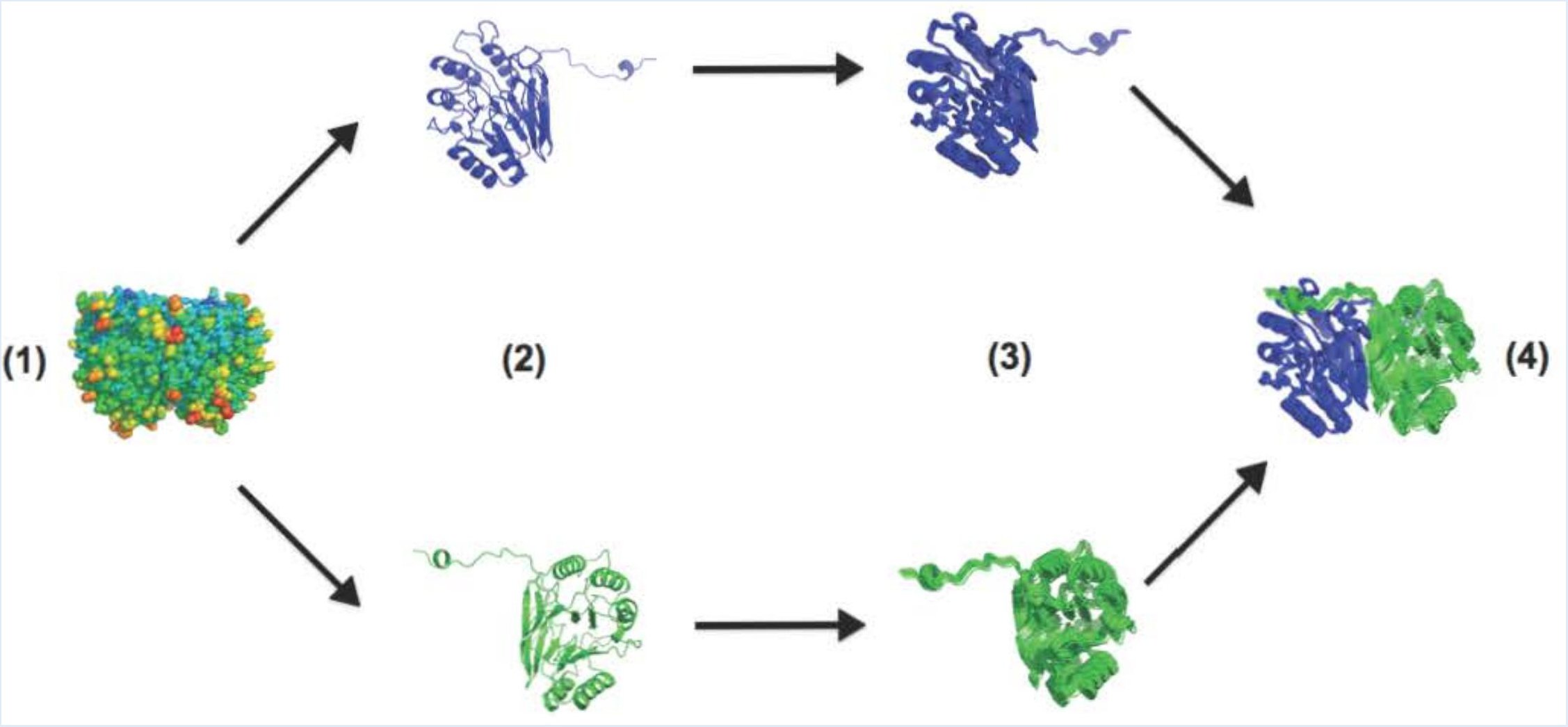
Overview of *Phenix.tls_as_xyz*. The input PDB (1) is broken down into its constituent TLS groups (2) and TLS ensembles are generated for each group independently (3). These groups are then re-assembled into the complete protein structure on a model-by-model basis (4).

### 2.3. phenix.diffus

*Phenix.diffuse* implements Guinier’s description of diffuse scattering (Guinier, 1963) **(Figure 3a)**. Diffuse scattering is calculated entirely from a series of unit cell “snapshots” contained in a multi-model PDB ensemble and assumes no motional correlation between crystal unit cells. This simplification ignores sources of disorder spanning multiple unit cells, which can contribute to experimentally measured diffuse scattering (Clarage *et al.*, 1992; Doucet & Benoit, 1987; Wall *et al.*, 1997). If desired, *phenix.diffuse* could model these large-scale effects through the analysis of a “supercell” containing multiple unit cell copies as implemented in several recent MD simulations of small proteins (Janowski *et al.*, 2013; Kuzmanic *et al.*, 2014). Guinier’s equation can be applied to arbitrarily sized crystalline regions; thus, a system of multiple unit cells allows for analysis of motions that occur between and across unit cells. In line with previous diffuse scattering simulations (Wall *et al.*, 2014), our program calculates structure factors for each ensemble member at the Bragg lattice positions, from which each term in Guinier’s equation is determined.

**Figure 3).**
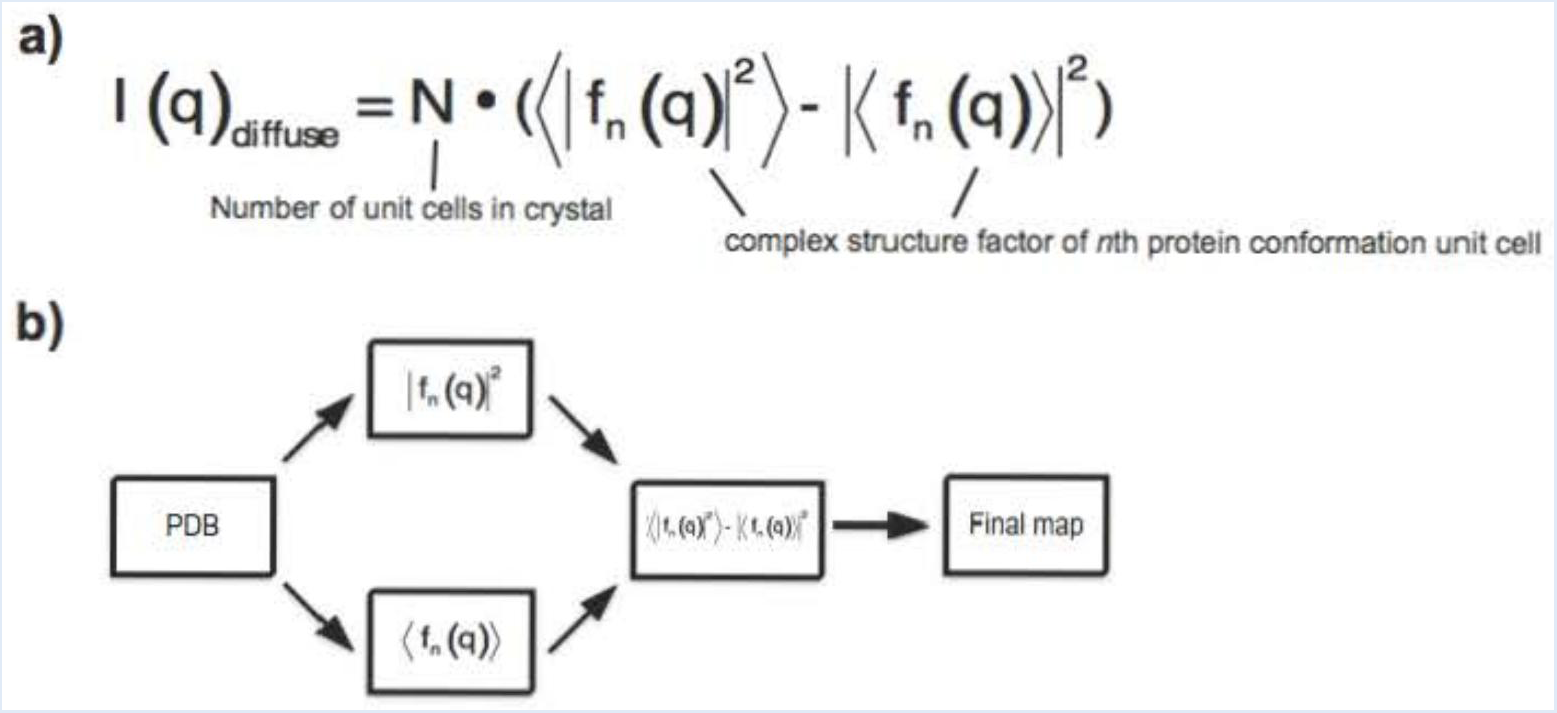
Overview of *Phenix.diffuse*. **a)** The general form of Guinier’s equation, Themotion to be analyzed is captured in a series of “snapshots” defined by the the multi-model PDB. **b)** The general program flow. Each term in Guinier’s equation is calculated separately from the structural ensembles and then combined to obtain the final map.

### 2.4. GpdQ TLS diffuse scattering simulation.

We simulated the diffuse scattering of each of the GpdQ TLS ensembles to 3.0Å resolution. Since the diffraction data for GpdQ PDB entry 2DXN extends to 2.9Å, our simulation should be sufficient for future comparisons with experimental maps. As the resulting diffuse scattering data is identical in format to descriptions of Bragg X-ray reflections, *phenix.reflection_statistics* was used to perform all statistical analyses. All reported correlation values are global Pearson correlation coefficients calculated between the described two sets of diffuse intensities. As previously mentioned (and described in Wall *et al.* (1997)), spherically symmetric sources of diffuse scattering contribute significantly to the observed intensity. In order to remove these confounding effects, we used the LUNUS software package (Wall, 2009) to subtract from each point the average radial diffuse intensity (**Supplemental Figure 2).**

### 2.5. GpdQ diffraction image processing and radial averaging.

Diffraction images used to determine the GpdQ Bragg structure were collected at the Advanced Photon Source (Lemont, IL) under cryogenic temperatures with 0.25^o^ oscillation wedges (Jackson *et al.*, 2006). Subsequent processing was performed using LUNUS (Wall, 2009). Pixels correlating to the beamstop shadow and CCD detector panels were removed with the LUNUS *punchim* and *thrshim* routines. Solid-angle normalization and beam polarization were corrected using *polarim* and *normim*. Mode filtering was applied as previously described (Wall *et al.*, 1997). The radial intensity profile was calculated from a single image using the *avgrim* function, which calculates radial intensities on a per-pixel scale.

## 3. Results

### 3.1. Diffuse scattering is dependent on TLS grouping.

The raw diffuse intensity predicted by each TLS refinement strategy rises as a function of the number of TLS groups **(Figure 4)**. The “entire molecule” and “monomer” maps show a similar range of intensity values: 0-4.52x10^6^ and 0-8.34x10^6^ respectively. The “subdomain” map displays a much wider dynamic range (0-4.71x10^8^) **(Supplemental Figure 1c)**. This trend likely results from an increase in the amplitude of TLS motion, particularly within the dimerization region of the “subdomain” model. **(Supplemental Figure 1)**. However, “sub-domain” map intensities greater than 1x10^7^ are limited to a resolution range of 11Å and lower. The “entire molecule” and “monomer” maps also possess “primary diffuse shell” regions surrounding the origin, though they only extend out to a resolution range of 30Å. This region will be particularly difficult to measure experimentally given the presence of a beamstop, which blocks access to signal around F_000_ (Lang *et al.*, 2014). Each diffuse map has a dip in radial intensity between the primary diffuse shell before the diffuse intensity rises in a second shell **(Figure 5a).** In contrast to the “sub-domain” map, the strongest diffuse intensities for the “entire molecule” and “monomer” maps occur within this secondary shell. The width between the primary and secondary diffuse shells decreases as the number of TLS groups increases, due to an expansion in the primary diffuse shell radius. As X-ray detectors can easily measure intensities in the regions of reciprocal space occupied by the secondary shell, a significant fraction of the diffuse scattering predicted by TLS refinement can potentially be compared to experimental data.

**Figure 4).**
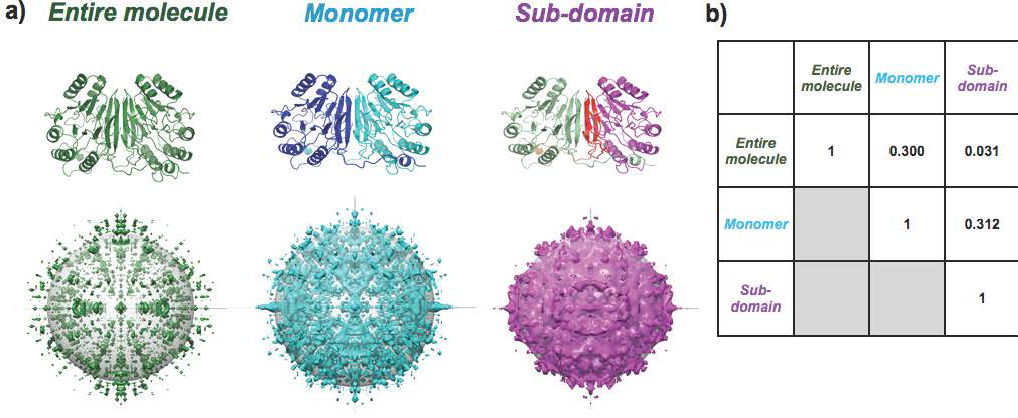
Differing TLS groups produce unique diffuse scattering. **a)** The GpdQ TLS groups projected onto the structure, along with the calculated diffuse scattering (looking down the L axis; grey sphere denotes 4Å resolution). The “Monomer” and “Subdomain” maps are shown at equivalent density thresholds, while “Entire molecule” is set at 60% of the density threshold. **b)** Pearson correlation coefficients between anisotropic maps.

**Figure 5).**
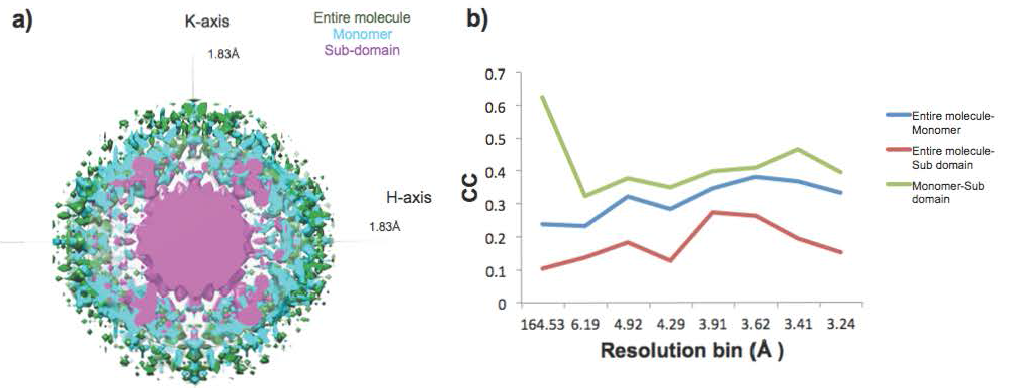
Comparison of simulated GpdQ TLS diffuse scattering maps. **a)** Cross-section of simulated TLS diffuse scattering maps. Primary and secondary diffuse intensity shells, separated by a gap, can be observed in each model. As the number of TLS groups increase, the intensity shells grow closer, predominantly due to an expansion in primary intensity shell size. **b)** Pearson correlation values between each set of maps across resolution bins.

To determine if the different TLS groupings yielded distinct diffuse scattering predictions, we calculated global Pearson correlation coefficients between each refinement’s anisotropic signal. The anisotropic comparison revealed little similarity between maps (CC range from 0.031 to 0.312) **(Figure 3)**. Comparing the correlation values across resolution bins reveals the anisotropic diffuse signal correlations remain consistently poor across scattering vector length **(Figure 5c)**. The large discrepancy between the maps calculated with different TLS models contrasts with the high similarity of experimental maps of anisotropic diffuse signal from different crystals of *Staphylococcal* nuclease (CC = 0.93) (Wall *et al.*, 1997). This result suggests the experimentally measured diffuse signal will be sufficiently precise to distinguish between TLS-related diffuse scattering models (Wall *et al.*, 2014).

### 3.2. Correlations between TLS groups can be detected by diffuse scattering.

Although TLS refinement assumes that each group moves as an independent rigid body, diffuse scattering can test whether there are correlated fluctuations between TLS groups. To illustrate this concept, we sequentially sampled the motions along each predicted translation and libration vector in opposing “parallel” and “antiparallel” directions from the “monomer” GpdQ TLS refinement **(Figure 6).** In contrast to the simulation in **Figure 4a**, here we have introduced correlated motion between GpdQ monomers. Next, we simulated the diffuse scattering produced by the “parallel” and “antiparallel” correlated motions. Both raw maps display strong secondary shell characteristics in combination with a weak primary shell of diffuse scattering **(Figure 6c).** A diffuse intensity difference map **(Figure 6d)** shows that discrepancies between the raw maps occur across the entirety of reciprocal space. Comparing the anisotropic diffuse intensity correlation across resolution bins reveals a general decreasing trend as the scattering vector length increases **(Figure 6e)**. In contrast to the previous TLS simulations, the correlation values are highest at low resolution. The low global Pearson correlation coefficient (0.375) demonstrates there are quantitative differences between the two maps. However, these intergroup correlation differences will be slightly more difficult to detect than changes between specific TLS models, where the correlation coefficients range from 0.031 to 0.312.

**Figure 6).**
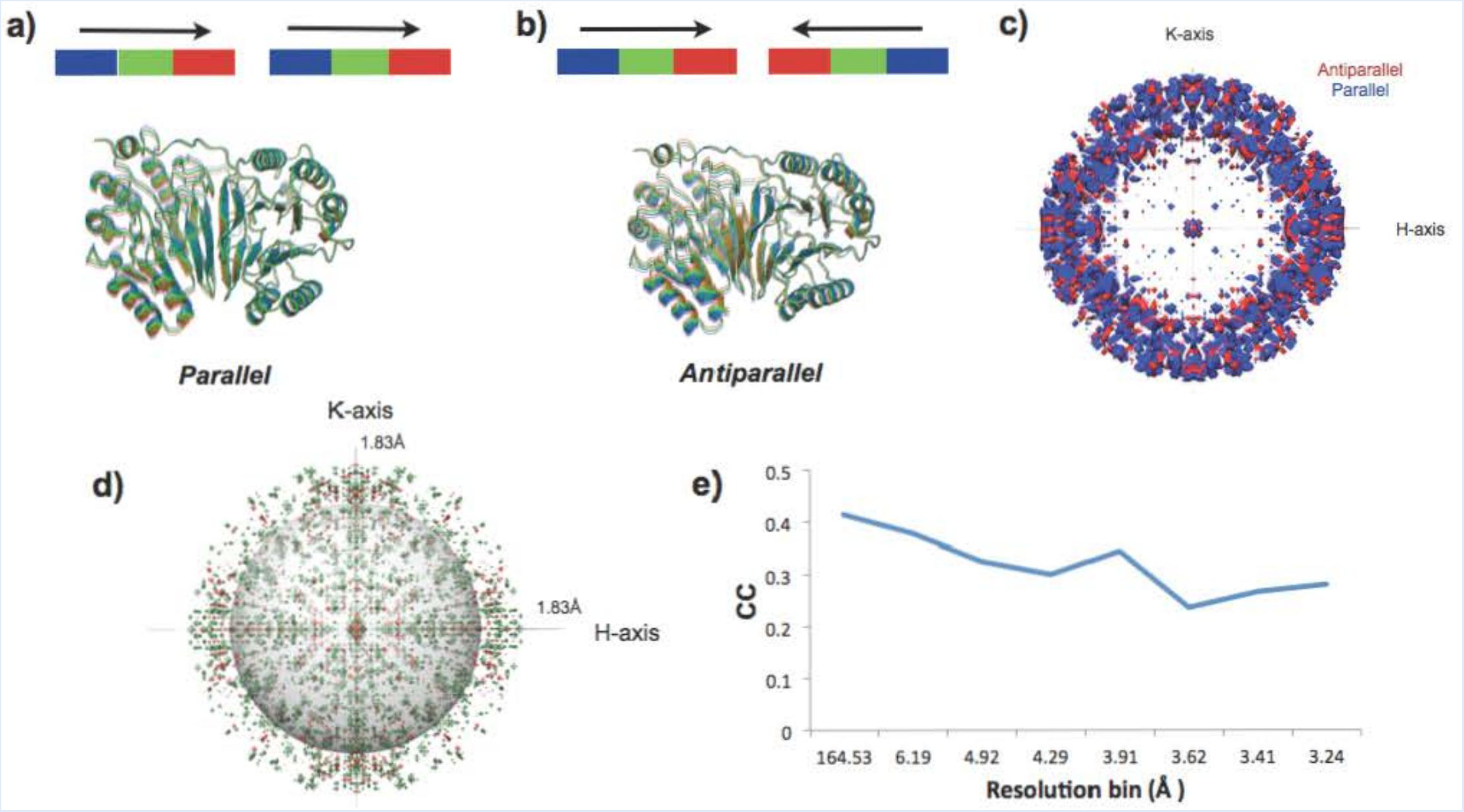
Different correlations between TLS groups produce unique diffuse scattering. Parallel **(a)** and antiparallel **(b)** TLS motions in GpdQ chains result in measurable differences between diffuse scattering patterns (CC = 0.375). Color bars indicate the directionality of the TLS motions; each color represents a unique molecular position. **c)** A map cutaway reveals strong secondary shell features with a small primary diffuse shell (looking down the L axis; grey sphere denotes 4Å resolution). **d)** Intensity differences between raw “anti” and “parallel” diffuse maps (green: positive, red: negative) highlights the qualitative changes caused by alternative TLS group correlations. **e)** Correlation values across anisotropic map resolution bins reveal highest correlation occurs between the maps at low resolution and decreases as a function of scattering vector length.

### 3.3. TLS models yield unique radial profiles of diffuse intensity.

We calculated the radial diffuse intensity profile for a GpdQ diffraction frame and for the three TLS refinements **(Figure 7)**. Although radial averaging removes the rich directional information present in diffuse scattering, this simplification has been successfully used to assess agreement between distinct diffuse maps (Meinhold & Smith, 2005; Meinhold & Smith, 2007). For the experimental GpdQ map, a peak at 8.5Å and a shoulder at 6Å are observed. None of these features are observed in the raw TLS radial profiles, except for a local maximum at 4.5Å and shoulder at 4Å for the “monomer” refinement. Rather, the dominating feature for each TLS simulation is the secondary diffuse scattering shell, which varies between maps in both width and maximum radial value. This result is not surprising, as the experimental diffuse scattering from GpdQ reflects a much broader group of correlated motions than simply TLS-related movement within the macromolecule. For example, disordered solvent is expected to significantly contribute to experimental diffuse measurements (Wall *et al.*, 1997). As solvent molecules were not modeled in our TLS ensembles, this is a likely source of the discrepancy between GpdQ experiment and simulation. The liquid-like motions (LLM) model, in which atoms interact only with nearest neighbors to produce a gelatinous crystalline environment, can also be used to explain the diffuse scattering intensity. Comparing the diffuse maps of *Staphylococcal* nuclease (Wall *et al.*, 1997), pig insulin (Caspar *et al.*, 1988) and hen egg-white lysozyme (Clarage *et al.*, 1992) to LLM models maximized correlations across distances of 6-10Å. Thus, a more thorough analysis involving several models of disorder must be applied to GpdQ to improve the fit to the experimental diffuse data.

**Figure 7).**
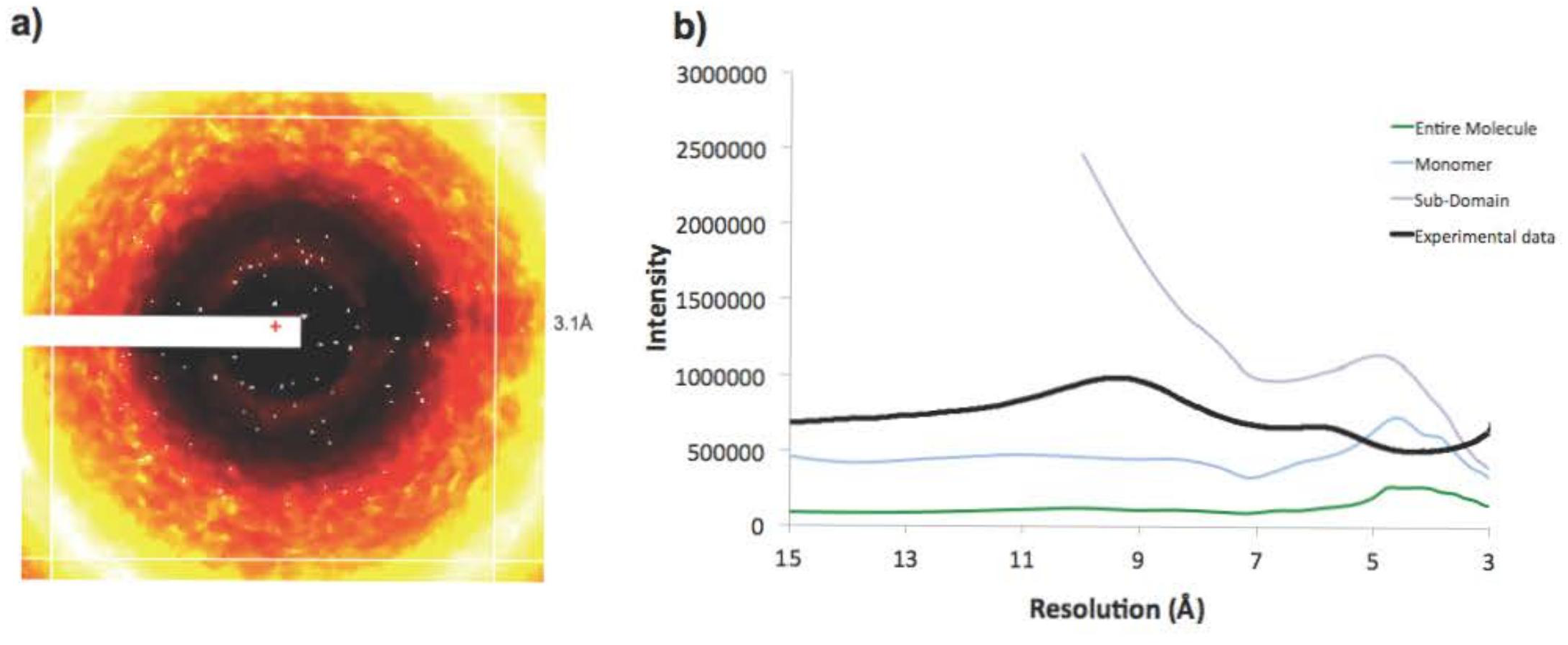
TLS models yield unique radial profiles of diffuse intensity. **a)** Mode filtered GpdQ diffraction image used for radial intensity calculation. The white regions correspond to pixels thrown out due to detector panel and beamstop artifacts, as well as Bragg scattering contamination. **b)** Radial diffuse intensity profiles for experimental and simulated GpdQ data. Resolution data below 15Å (roughly corresponding to the primary diffuse shell) were removed for more accurate visual comparison. The “Subdomain” map exceeds the limits of the Y-axis at lower than 10Å resolution.

### 3.4. Distinct patterns of diffuse signal can be calculated at Non-Bragg Indices

While *phenix.diffuse* currently calculates the diffuse signal under Bragg peaks, diffuse scattering occurs throughout the entirety of reciprocal space. To more completely sample reciprocal space between the Bragg spots, we increased the unit cell boundaries. Expanding the unit cell in real space allows for finer sampling of the underlying Fourier transform **(Figure 8)**. The resulting structure factors can be re-scaled to the original lattice points, leading to fractional *hkl* sampling. These fractional values are then assigned to the nearest integer *hkl* index and averaged, leading to a single diffuse intensity value associated with each Bragg peak. Although it is clearly possible to output a map consisting of these fractional values and thereby produce a more accurate picture of diffuse scattering, we chose the integer values because diffuse scattering processing techniques commonly calculate the average diffuse intensity across pixels within a 1x1x1 voxel around each Bragg point (Wall, 1996). This average value is then assigned to the *hkl* index, leading to the same 1:1 correlation between lattice points and diffuse intensity values. Although it is tempting to use this method in our current analysis, the unit cell expansion method does not maintain the expected crystallographic symmetry for any crystal system with a screw axis. Introducing vacuum into our structure factor calculations will satisfy other symmetry operations, but as GpdQ possesses a screw axis we are currently unable to more finely sample its predicted diffuse scattering. Therefore, we can use this method to compare data between simulated models of motion, but not between simulated models and experimental data. More advanced simulation methods will need to incorporate screw axes, either by defining a new supercell for simulation or directly calculating structure factors at fractional *hkl* indices. Cognizant of these limitations, we calculated the diffuse scattering of each of the GpdQ TLS ensembles to 3.0Å resolution in a P1 cell, with a sub-sampling of 4x4x4 around each Bragg lattice point (**Figure 8c).** These calculations confirm that each TLS motion produces distinct patterns of diffuse signal throughout reciprocal space.

**Figure 8).**
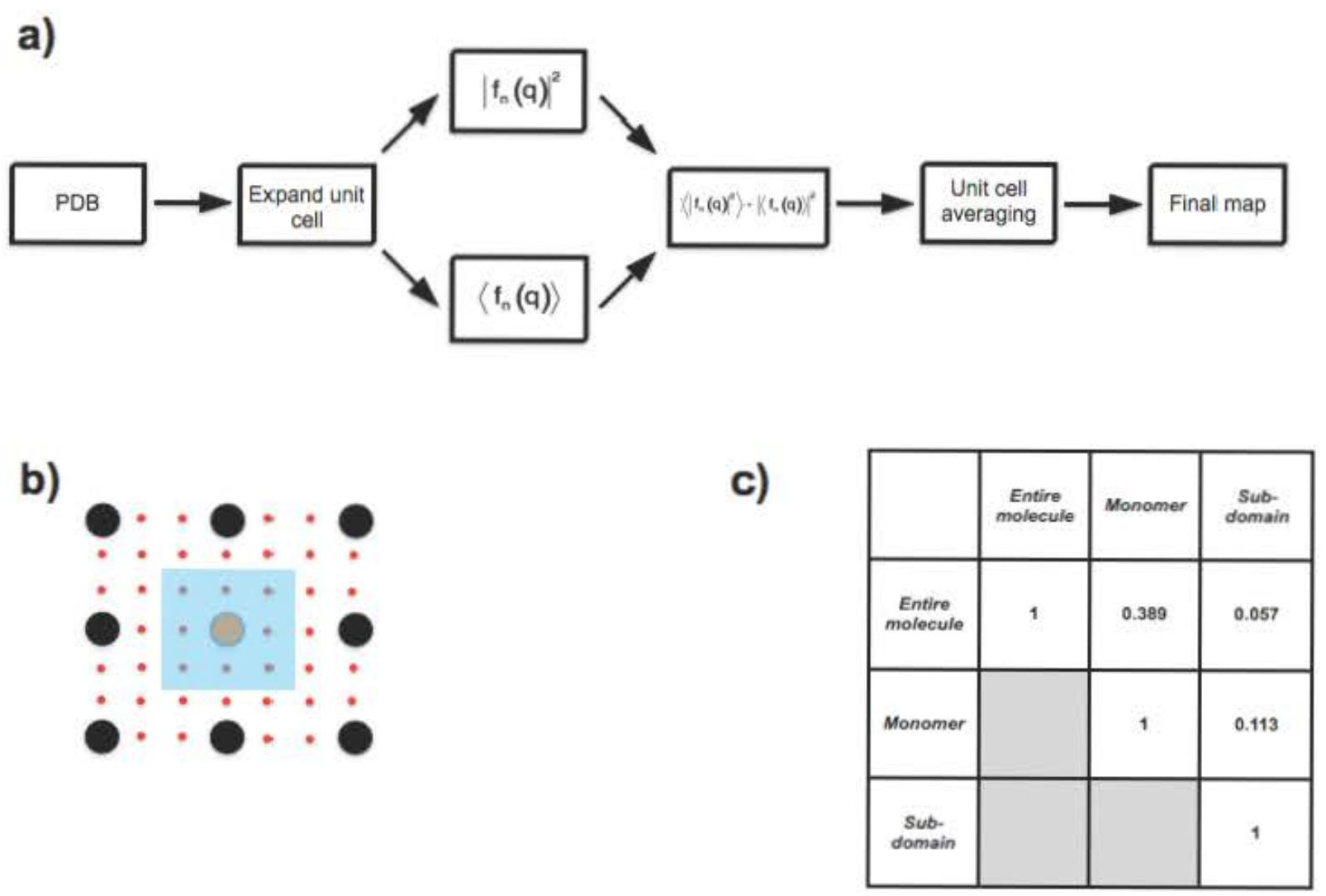
Unit cell expansion allows for reciprocal space subsampling. **a)** The inputPDB’s unit cell is expanded to create the desired unit cell sampling, each term in Guinier’s equation is calculated separately and then the second term is subtracted from the first to obtain the diffuse intensity. The “pseudo-unit cells” are then averaged across, producing the final diffuse scattering map. **b)** Unit cell expansion allowing for 3x subsampling of reciprocal space. True/”pseudo” Bragg peaks are shown in black/orange and red, respectively. The intensity values of the eight pseudo peaks and one orange peak in the blue box are averaged and the resulting value is assigned to the orange peak’s Bragg index. **c)** Pearson correlation coefficients between maps.

## 4. Discussion

Accurate modeling of conformational dynamics is important for understanding macromolecular function. Although many models may fit the existing data equally well, they can often suggest different correlated motions. Our results suggest that comparisons to experimental diffuse scattering can break the degeneracy between different TLS refinements, as each TLS motion produces markedly different diffuse patterns. For example, alternative correlations between TLS groups have equivalent average electron density, but result in unique diffuse scattering predictions. More generally, any model proposed through TLS refinement should agree with the experimental diffuse data, as this data directly reflects the existing protein disorder (Moore, 2009).

Despite this synergy between TLS refinement and diffuse scattering, there are many potential complications when applying TLS X-ray refinement to model protein dynamics. As the T and L matrices describe independent translations and librations, these motions must be physically sensible. Our review of protein structures deposited in the Protein Data Bank indicates that 88% of refinements employing TLS (24% of the total PDB) do not satisfy this physical requirement (Urzhumtsev *et al.*, complementary paper). We hypothesize that this discrepancy arises due to a lack of restraints applied to refined TLS parameters to ensure their physical plausibility. Even if this criterion is met, current TLS refinement methods still do not impose chemical restraints between TLS groups, which can lead to displacements that are chemically unreasonable. Our TLS refinement of the GpdQ subdomain is one such example, as it produces rigid body displacements that extend across the entirety of the unit cell **(Supplemental Figure 1c)**. Thus, validation checks of TLS refinement (such as those implemented in *phenix.tls_models*) are critical, as is employing TLS refinement within a broader framework of restraints. Alternative techniques, such as the Phase Integrated Method (PIM), which derives anisotropic B factors from low-frequency normal modes (Chen *et al.*, 2010), may significantly improve the biochemical accuracy of modeling efforts. In PIM, the fit between model and experiment is significantly improved by calculating normal modes in the context of the asymmetric unit rather than individual molecules (Lu & Ma, 2013).

With the increasing availability of modeling tools such as *phenix.diffuse*, the lack of high-quality three-dimensional datasets is now a key bottleneck in diffuse scattering analysis. One challenge in data collection is that long X-ray exposures can be required to reveal diffuse features. This can lead to blooming around saturated Bragg spots in diffraction images collected using commercially-available charge-coupled device (CCD) area detectors (Gruner *et al.*, 2002) Blooming can artificially increase pixel values between the Bragg spots, where the diffuse intensity is measured (Glover *et al.*, 1991). Although CCD detectors can be configured to eliminate spot blooming at the cost of decreasing dynamic range (Wall, 1996; Wall *et al.*, 1997), this configuration is not available in commercial detectors. The development of pixel array detectors, which possess much higher dynamic ranges as well as very small point-spread functions, has opened the door to more accurate measurement of diffuse signal. Additionally, methods for processing diffuse scattering data from raw image frames to complete reciprocal space map are under active development (Wall *et al.*, 2014). These methods will be applied to new datasets of simultaneous Bragg and diffuse scattering data. Instead of being included in the background corrections in estimating Bragg peak intensities, these diffuse intensities will increase the data available for refinement, enable more accurate quantification of interatomic distances (Kuzmanic *et al.*, 2011), and allow for the simultaneous refinement of multiple coupled protein motions (Wilson, 2013).

## Acknowledgements

J. S. F. is a Searle Scholar, a Pew Scholar, and a Packard Fellow. Work in the lab of J. S. F. is supported by NIH OD009180, GM110580, and NSF STC-1231306. P. D. A., P. V. A., and T. C. T are supported by NIH grant GM063210. N. K. S. was supported by NIH grant GM095887. A. U.thanks the French Infrastructure for Integrated Structural Biology (FRISBI) ANR-10-INSB-05- 01 and Instruct as part of the European Strategy Forum on Research Infrastructures (ESFRI). M. E. W. is supported by the US Department of Energy through the Laboratory-Directed Research and Development program at Los Alamos National Laboratory. This work was supported by the Program Breakthrough Biomedical Research, which is partially funded by the Sandler Foundation.

---

**Supplemental Table 1).** Eigenvalues of GpdQ TLS refinement matrices. **a)** “Entiremolecule” **b)** “Monomer” **c)** “Sub-domain”. It is important to note that, for the “Subdomain” refinement, L#5 and L#6 have negative eigenvalues. Due to their low value, however, these eigenvalues were set to zero for subsequent calculations.

| T | L |
| --- | --- |
| 0.854 | 1.405 |
| 0.258 | 0.717 |
| 0.338 | 0.172 |

*Supplemental Table 1b)*
| T<br>Monomer A | L<br>Monomer A | T<br>Monomer B | L<br>Monomer B |
| --- | --- | --- | --- |
| 0.873 | 1.843 | 0.850 | 1.896 |
| 0.236 | 0.021 | 0.192 | 1.103 |
| 0.327 | 0.822 | 0.329 | 0.500 |

*Supplemental tables 1c)*
| T<br>Sandwich<br>(A) | L<br>Sandwich<br>(A) | T<br>Dimerization<br>(A) | L<br>Dimerization<br>(A) | T<br>Cap<br>(A) | L<br>Cap<br>(A) |
| --- | --- | --- | --- | --- | --- |
| 0.917 | 0.005 | 0.942 | 1.420 | 0.902 | 0.154 |
| 0.247 | 1.396 | 0.313 | 0.181 | 0.475 | 0.062 |
| 0.367 | 0.957 | 0.375 | 0.057 | 0.323 | 0.005 |

| T<br>Sandwich<br>(B) | L<br>Sandwich<br>(B) | T<br>Dimerization<br>(B) | L<br>Dimerization<br>(B) | T<br>Cap<br>(A) | L<br>Cap<br>(A) |
| --- | --- | --- | --- | --- | --- |
| 0.940 | 0.216 | 0.897 | 1.07 | 0.938 | 0.155 |
| 0.170 | 1.265 | 0.267 | 0.059 | 0.638 | **-0.003** |
| 0.399 | 0.838 | 0.368 | **-0.001** | 0.477 | 0.031 |

**Supplemental Table 2).** Multi-model ensembles are necessary for adequate random sampling of TLS motions. Two ensembles independently sampling the underlying TLS distributions were used to generate anisotropic diffuse scattering maps. Global CC values between the two maps are shown. These simulations were conducted in triplicate, producing the CC standard deviation shown in parentheses. All maps were simulated to 3 Angstrom resolution.

|  | **10** | **50** | **100** | **500** | **1000** |
| --- | --- | --- | --- | --- | --- |
| **Entire**<br>**Molecule** | 0.886<br>(0.027) | 0.956<br>(0.019) | 0.988<br>(0.005) | 0.996<br>(0.002) | 0.999<br>(0.000) |
| **Monomer** | 0.809<br>(0.087) | 0.924<br>(0.008) | 0.952<br>(0.005) | 0.992<br>(0.002) | 0.997<br>(0.001) |
| **Sub-**<br>**domain** | 0.944<br>(0.012) | 0.984<br>(0.005) | 0.992<br>(0.001) | 0.999<br>(0.000) | 0.999<br>(0.000) |

**Supplemental Figure 1).**
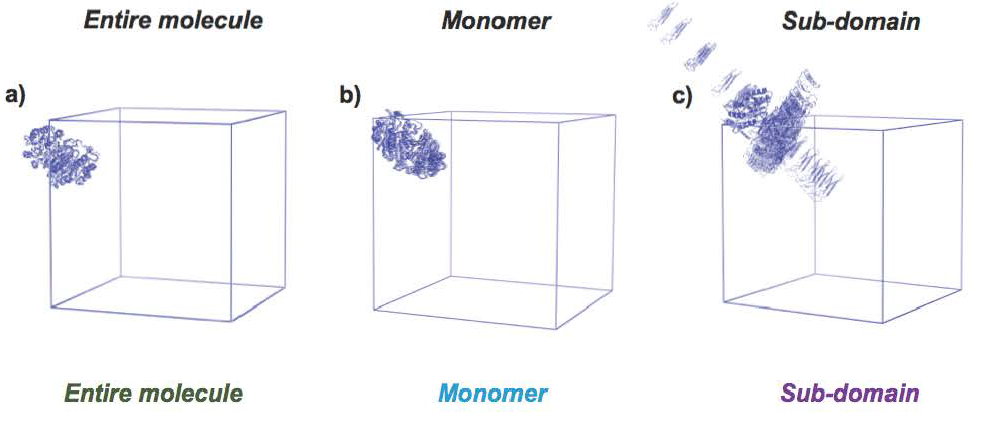
Structural ensembles of GpdQ TLS motions. Each TLS PDB ensemble is shown as a single asymmetric unit outlined by the unit cell. An increase in overall motion is apparent going from left to right. The 20 member ensemble is shown for visual simplicity. It is important to note that the chemically unreasonable motion produced by the sub-domain TLS refinement is not immediately apparent from the T and L eigenvalues presented in Supplemental Table 1, highlighting the need for the more thorough matrix analysis presented in our accompanying paper.

**Supplemental Figure 2).**
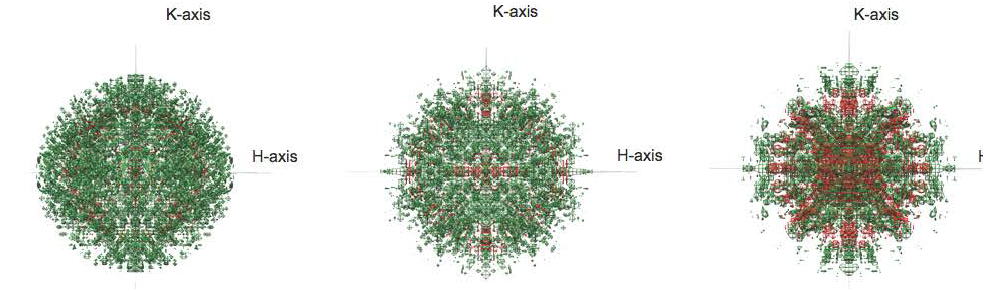
Anisotropic diffuse scattering maps. Positive and negative anisotropic density is shown as green and red mesh, respectively. Absolute threshold levels shown for the positive and negative signals are equivalent. The maps are shown to their full 3 Angstrom resolution limit.

